# MDCompress: better, faster compression of molecular dynamics simulation trajectories

**DOI:** 10.64898/2025.12.19.695527

**Authors:** Marek Kokot, Amitava Roy, Travis J Wheeler, Sebastian Deorowicz

## Abstract

**Motivation:** Molecular dynamics (MD) simulations model the physical movements of atoms in biomolecular systems over time, providing atomic-resolution insight into conformational changes, binding events, and dynamic behaviors that cannot be captured by static structures alone. As such, MD simulations are playing an increasingly important role in understanding the functional roles and molecular interactions of proteins. However, trajectories from these simulations can be extremely large, often reaching tens of gigabytes for a single simulation of modest duration. This creates substantial challenges for storage and data transfer, motivating efficient compression strategies. Furthermore, many downstream analyses require extraction of only a subset of frames or specific atoms from the full trajectory, so an ideal compression format should support rapid random-access decompression of such samplings without requiring full file decompression.

**Results:** Here, we introduce MDCompress, a new trajectory compression format and accompanying software implementation that meets these goals. MDCompress produces compressed trajectory files that are 15–37% smaller than those generated by the widely-used XTC format, while achieving faster compression and decompression speeds through efficient multithreading.

**Availability and Implementation:** The MDCompress software and library are released under an open license (BSD-3) and may be downloaded at https://github.com/refresh-bio/mdcompress

**Supplementary information:** Supplementary data are available online.

## Introduction

The structure of a protein informs its function in broad strokes, providing insight into its capacity to bind other molecules, catalyze reactions, act as a structural component, transduce signals, transport and store ligands, and more. Recombination of structural domains, together with subtle modifications within those domains, has generated the vast diversity of form and function observed across the proteome. Determining protein structures has been the focus of decades of effort, historically dominated by X-ray crystallography and supplemented by cryo-electron microscopy (cryo-EM), nuclear magnetic resonance (NMR) spectroscopy, and other methods. Since 1971, experimentally determined structures have been catalogued in the Protein Data Bank (PDB [1]), enabling researchers to share new discoveries and explore previously solved structures. The impact of this archive (now encompassing over 200,000 entries) cannot be overstated. For example, PDB data served as the foundation for deep neural-network models that, as of 2020, began predicting high-accuracy structures for arbitrary proteins, triggering a paradigm shift in the way that researchers consider understudied or novel proteins.

Thanks to advances in structure-prediction algorithms, massive repositories of predicted protein models now exist. The AlphaFold Protein Structure Database [2] contains approximately 214 million predicted structures at the time of this writing, and the ESM Metagenomic Atlas [3] catalogs around 772 million. Because each structure file encodes the three-dimensional coordinates of every atom, these databases become extremely large – AlphaFold’s repository alone approaches 23 TiB. To lower the storage and handling burden of such vast collections, we recently developed a software tool (ProteStAr [4]) that achieves high-throughput, lossless compression of protein structures. ProteStAr works by converting absolute atomic coordinates into a reduced coordinate frame via prediction of the atom coordinates using a reference model, applying coordinate quantization, and then using an entropy-coder optimized for three-dimensional spatial data. This pipeline yields compression ratios substantially better than general-purpose compressors, while preserving exact geometry for downstream analyses.

Importantly, a single snapshot of a protein structure, whether obtained by crystallography, cryo-EM, NMR, or AI-based prediction, does not capture the dynamic behavior that underlies function. Proteins and their interacting partners are flexible molecules whose conformational fluctuations regulate the feasibility, kinetics, and outcomes of interactions. For example:

- Population shift in small-molecule binding: Many enzymes populate an ensemble of conformations that differ in loop orientation, side chain rotamers, and even subtle backbone adjustments, and continually fluctuate between them. A small-molecule ligand (a substrate, activator, or inhibitor) preferentially interacts with a subset of these pre-existing conformations, shifting the dynamic equilibrium from conformers with weak ligand-binding affinity (“open” states) toward conformers with stronger ligand-binding affinity (“closed” states), and thus altering activity. These dynamic fluctuations between “open” and “closed” states directly impact function.
- Allostery: For many proteins, an event at one site on a protein influences structure and function at a distant site. This process, called allostery, is a central mechanism for signal transmission in many receptors and enzymes. For example, in G-protein–coupled receptors (GPCRs), ligand binding in an extracellular pocket drives concerted rearrangements of the transmembrane helices that are propagated to the intracellular face, creating a G-protein binding interface and initiating downstream signaling.
- Intrinsically disordered regions: Many proteins (or regions within proteins) lack a single well-defined fold and instead sample broad conformational ensembles in solution. These intrinsically disordered regions (IDRs) can adopt structure when interacting with binding partners.
- Macromolecular assemblies: Macromolecular assemblies are multi-component machines whose function requires components to assemble in a specific sequence and then undergo coordinated large-scale rearrangements. For example, in the ribosome, the large and small subunits rotate relative to each other to translocate tRNA and mRNA, while dynamic conformational changes in ribosomal proteins and rRNA govern the peptidyl-transferase reaction.

Together, these examples illustrate that a static structure provides only a partial picture of the function of a protein. To uncover the mechanistic underpinnings of molecular recognition, catalysis, or signal transduction, one must consider the ensemble of conformations and the transitions linking them.

Recording protein dynamics at full atomistic resolution remains beyond current experimental capabilities. However, physics-based molecular dynamics (MD) simulations have become indispensable in structural biology, driven by methodological advances and the widespread availability of GPUs. Just as the static structures in the Protein Data Bank (PDB) enabled the development of transformational AI models for structure prediction, the global corpus of MD trajectories holds immense promise as training data for next-generation AI methods. Such models have the potential to revolutionize research tied to protein-protein interactions, molecular transport, enzyme engineering, functional genomics, drug discovery, toxicity, bioremediation, immunology, and more. This fact has motivated a recent push to create [5, 6, 7, 8, 9] or gather [10] MD simulations of many thousands of systems (including our own repository for community-contributed simulations, MDRepo [11]).

### MD simulation file size and compression

Today, it is common to produce a simulation of a system with tens or hundreds of thousands of atoms over a duration of many 10s of nanoseconds. While conventional MD simulations typically advance in ~2 femtosecond time-step increments, the resulting full-system coordinates are stored at larger time steps, often in increments of 1–10 picoseconds. This sampling allows capture of conformational diversity across nanosecond-scale molecular processes, while somewhat limiting storage requirements. Even with this temporally sparse coordinate sampling, MD simulation files can be quite large, often reaching tens of gigabytes even for a single simulation of modest duration and molecule size, so that the total collection of accessible MD data will soon be measured in petabytes. The challenges and costs of storing and transporting such large data sets motivate development of new methods for compressing protein dynamics trajectory files.

A simple way to store an MD trajectory is to capture time-ordered frames, where each frame contains the x/y/z coordinates of all atoms in sequence. The landscape of conceptually similar uncompressed formats is fragmented across MD simulation tools, with common options including TRR (GROMACS [12]), NetCDF (AMBER [13]), and DCD (CHARMM [14], NAMD [15], ACEMD [16]). Either as native output or via format conversion, it is common to store trajectories in the compressed XTC format. Note that XTC format stores only the coordinates of the trajectory, and does not include other features often optionally stored in uncompressed formats, such as per-atom velocities and forces. The XTC compression algorithm is “lossy” (it discards precision below the assigned rounding threshold), but retained precision is generally well within chemical-accuracy tolerances. All of the above formats are interconvertible using libraries such as MDTraj [17].

XTC files are generally ~3x smaller than uncompressed coordinate-only formats, and thus are a common format for storing large collections of simulations (including in our repository, MDRepo). One substantial weakness of the XTC format is the lack of random access capability: because each frame is block-compressed and there is no top-level index, it is not possible to index directly into the file to find coordinates for a specified set of frames or atoms. As a result, extraction of subsets of data from an XTC-formatted file requires full file traversal. This limitation will become increasingly relevant as researchers develop AI models for predicting protein dynamics and interactions – such models will generally be trained on down-sampled data from trajectories, and efficient extraction will be at a premium.

Motivated by the goals described above, we have developed a new format for compressing molecular dynamics simulations, along with a reference software implementation. The software, MDCompress, results in compressed files that are smaller than XTC-formatted files, with fast multi-threaded compression/decompression performance and support for random access extraction on a per-frame or per-atom basis.

### A history of MD simulation compression

The XTC format reduces trajectory size by combining three techniques. First, coordinates are quantized to a user-chosen precision (for example, rounding to the nearest 1 pm), converting floating-point positions to small integers and introducing a bounded, configurable loss. Second, XTC records delta values between successive frames rather than full absolute positions; those deltas tend to be small and are stored using bit-packing (the minimum number of bits required per value), which dramatically reduces raw size. Finally, each frame’s packed delta block is compressed independently using zlib. Since its introduction in the mid-1990s, a few MD compression alternatives to XTC have been proposed over the years.

The TNG format [18] is block-structured with explicit indexing, allowing for frame-level random access. It combines quantization, predictive coding, and entropy compression, and metadata support to achieve both higher compression ratios, while allowing either lossy or lossless storage. PMC [19] incorporates explicit bond connectivity into its coordinate prediction model, using information about linkage between atoms to guide the reconstruction of atomic positions. This bond-aware representation can improve accuracy in cases where the molecular topology is well defined and stable, but reliance on a static topology limits its usefulness to systems with stable bonds and makes it unsuitable for cases where connectivity is absent or changes during the simulation. SZ3 [20] and MDZ [21] apply general-purpose scientific data compression techniques, such as predictor–quantizer–entropy pipelines, to atom coordinate arrays. These approaches can achieve strong error-bounded compression ratios, but the published implementations were primarily released as proofs of concept for evaluation, and with limited tooling and documentation, they remain difficult to adopt in practical MD workflows.

In this article, we introduce MDC, a novel MD trajectory compression format, together with a reference software implementation. MDC uses advanced prediction techniques for atom coordinates for improved compression and yields files that are smaller than XTC while enabling significantly faster compression and decompression. MDC organizes data into distinct components, called *containers* [4], that can be processed independently; this enables both efficient parallel processing and rapid random access queries of various types (e.g., all atoms in a single frame, complete trajectories for a few atoms). Currently, MDC allows metadata and atom coordinates to be stored, the same as for XTC. Moreover, it is an open format so that new container types can be added for velocities, forces, or energies. We provide a command-line application, called MDCompress, that supports various input types for compression and decompression; we also provide decompression libraries for a few popular programming languages.

## Methods

### Overview

The fundamental idea behind MDCompress is that (1) it is possible to quickly compute a reasonably-good prediction of the location of an atom, and (2) the difference between the predicted and true location of the atom can be stored in a reduced number of bits using entropy encoding (specifically, a range coder [22]). This is similar to the approach used in XTC compression, differing in the method used to predict atom positions. In the XTC format, each atom’s predicted current position is the same as its position in the previous frame; meanwhile, MDCompress introduces an advanced prediction scheme that yields much smaller prediction errors, which can therefore be stored with fewer bits. MDCompress also organizes computation in a manner that enables parallelism with good scaling properties.

### Data preparation

Atoms in a trajectory are grouped by the molecule they belong to (see Fig. 1a for example):

**Fig. 1.**
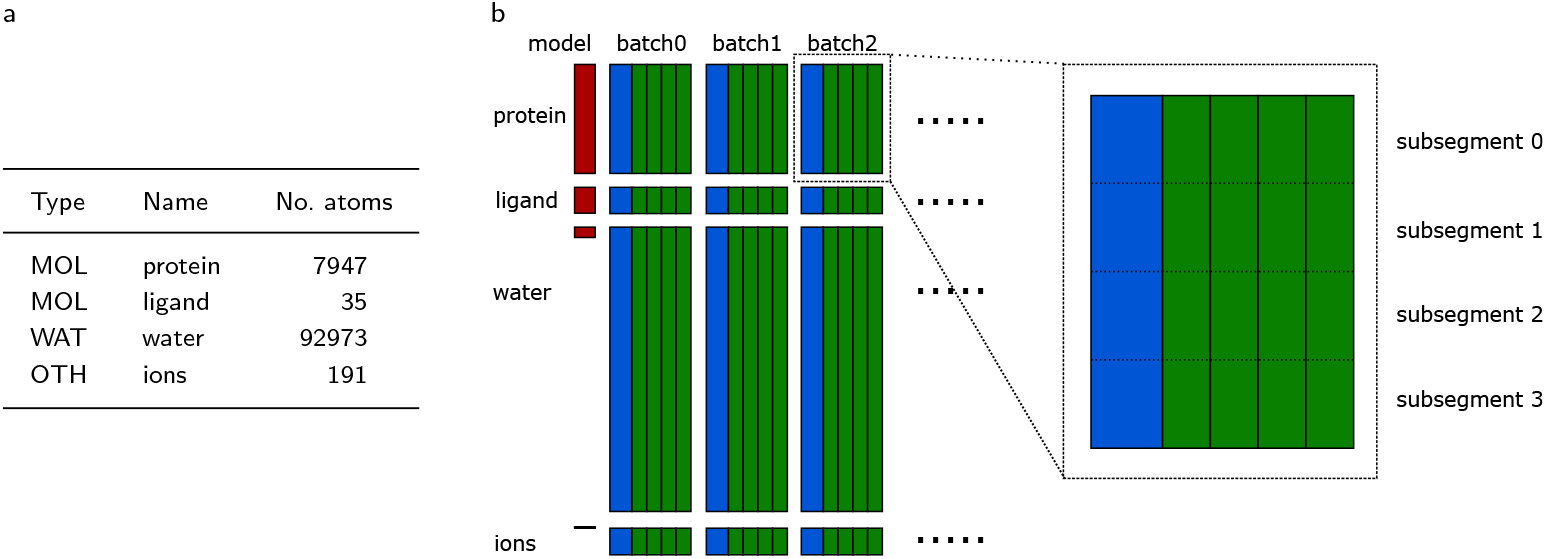
(a) Description of the segments for the full MDR00004434 trajectory, a simulation of a Nicotinamide phosphoribosyltransferase in complex with a ligand downloaded from MDRepo [11]. (b) Compressed archive contents: red blocks represent the model description, blue blocks represent anchor frames, and green blocks represent delta frames. The zoomed part shows the internal split of one segment into four subsegments.

- *MOL*—each molecule with at least six atoms (e.g., protein, ligands) is assigned to a separate *MOL* segment;
- *WAT* —water molecules that appear consecutively in the structure file are assigned to a single *WAT* segment; there may be multiple *WAT* segments;
- *OTH* —very small non-water molecules (including ions) that appear consecutively in the structure file are assigned to an *OTH* segment; there may be multiple *OTH* segments.

A trajectory is processed in *batches* of consecutive frames. Each batch of *n* frames starts with an *anchor* frame, followed by *n*−1 *delta* frames. The processing of each segment in each batch is independent. The compressed batches are stored in a container file (Fig. 1b). MDCompress offers five compression levels for various trade-offs between speed and compression ratio.

### Model construction

In the preprocessing stage, MDCompress analyzes a user-specified number of initial frames (default *m* = 100), and for each segment builds a *model* that will assist in predicting the positions of atoms. In each segment, atoms are considered in the order that they appear in the structure file. Let us use 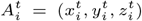 for the *i*-th atom’s coordinates in the *t*-th frame of the trajectory.

Position prediction is most complex in a *MOL* segment. For each atom *A*_*i*_, MDCompress considers the *b* (default 100) previous atoms as they appear in the structure file, seeking to identify three other atoms that are likely to best inform the prediction of the position of *A*_*i*_ throughout the trajectory. MDCompress considers atoms *A*_*j*_, such that max(1, *i* − *b*) ≤ *j* ≤ *i* − 1, because they are expected to be consistently close to *A*_*i*_ and are certain to have been computed by the time the prediction of *A*_*i*_ is computed. From this pool, MDCompress selects the three atoms that have the smallest average distance from *A*_*i*_ across all *m* initial frames: *A*_*p*(*i*)_, *A*_*q*(*i*)_, and *A*_*r*(*i*)_. These three *reference* atoms define a coordinate system for *A*_*i*_, and serve as the core of the *MOL* segment model; for each *A*_*i*_, the model also stores on whether the true position of is above the plane defined by its reference atoms.

For *WAT* segments, the model stores the median *d*_OH_ and *d*_HH_ distances (oxygen to hydrogen and hydrogen to hydrogen) for all water molecules in the initial frames. The model contains no information for *OTH* segments.

### Compression of segments

Compression is performed independently for each segment-batch pairing. For a given segment and batch, let 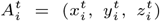 represent the coordinates of the *i*-th atom in the *t*-th frame of the batch (*t* = 0 for anchor frame). The compression of each atom’s coordinates starts by predicting their values 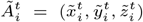, then encoding the prediction errors with a range coder. The decompressor mimics these estimations. Coordinates are processed sequentially, so, e.g., after encoding 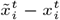 (prediction error for 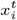), we can use 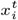 when predicting 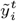, as 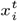 is known both for the compressor and decompressor.

#### Predicting *OTH* positions

The prediction of atom positions in *OTH* segments is the simplest. For anchor frames, the predicted position depends only on the true position of the preceding atom ( 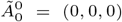 and 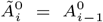) in delta frames, the predicted position of an atom depends on its true position in the previous frame 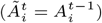

#### Predicting *WAT* positions

To predict atom positions in *WAT* segments, MDCompress treats each water molecule as a unit. For simplicity of presentation, let us assume that the *i*-th atom (*i* mod 3 = 0) is oxygen and the (*i* + 1)-th and (*i* + 2)-th are hydrogens of the same molecule. For oxygen atoms in the anchor frame, the predicted position is 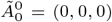 and 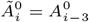, while 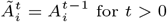 for *t >* 0. For the first hydrogen, the predicted x and y coordinates are 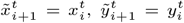 Given the oxygen position 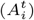 and these x and y coordinates, we can improve prediction accuracy for 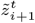 by observing that only two positions are consistent with those coordinates and the atomic distance *d*_OH_. MDCompress therefore computes the two candidates for 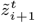 benefiting from the fact that one of them should be very close to 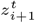 (Fig. 2a). MDCompress spends 1 bit) to encode which of the two candidates was used.

**Fig. 2.**
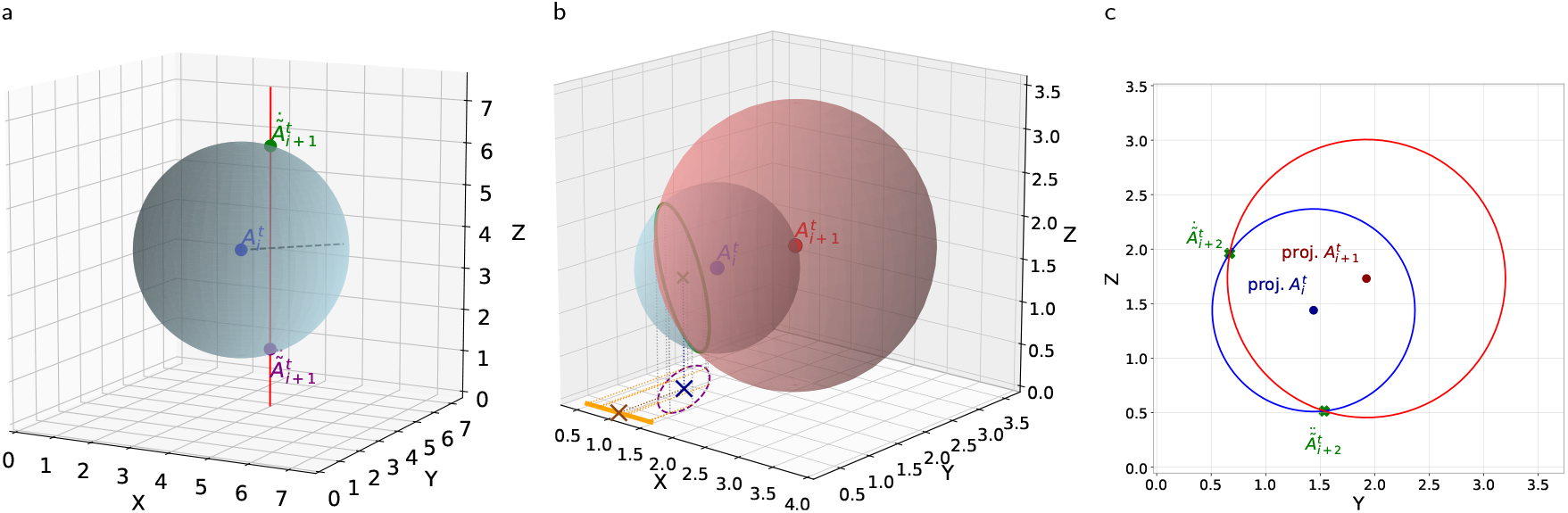
Illustration of the prediction of hydrogen atoms coordinates for water molecules: (a) Prediction of the first hydrogen, 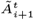. The blue point epresents the oxygen atom 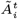 The radius of the sphere is *d*_OH_, the median distance of a hydrogen from its oxygen. After establishing the x and y coordinates of the hydrogen atom, we can draw a line, which intersects the sphere at two points – these are the candidates for the position of the first hydrogen atom. (b) Prediction of the second hydrogen atom, 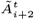. For 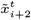, we consider two spheres centered at the positions of the oxygen (blue) and first hydrogen (red) atoms, with radii *d*_OH_ and *d*_HH_, respectively. The second hydrogen atom should be on the marked circle (intersection of the spheres). The projection of the center of the circle onto the X axis (brown cross) is the prediction of 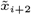. (c) Prediction of 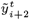. We again consider two spheres for oxygen and first hydrogen and intersect a plane 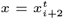 with these spheres. The red and blue points represent projections of the oxygen and first hydrogen atoms on this plane. The red and blue circles represent the intersection of the spheres and this plane. The intersections of those circles (green points) represent the predicted positions of the second hydrogen atom: 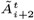

When predicting the position of the second hydrogen, additional compression is possible. In compression levels from 1 to 4, MDCompress predicts 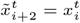 for the second hydrogen. In level 5, however, a more computationally intensive strategy is employed: two spheres are identified, one centered at the oxygen atom 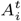 with radius *d*_OH_, the other centered at the first hydrogen atom 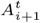 with radius *d*_HH_. The second hydrogen atom 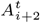 should be located somewhere on the intersection of the spheres, so we calculate the intersection (a circle) and project it onto the *z* = 0 plane, and then onto the X axis. The center of this segment is used as the predicted x-coordinate, 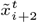 (Fig. 2b). In compression levels 1 and 2, the y-coordinate is predicted as 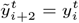 In higher levels, spheres are computed as above, based on 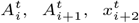 *d*_OH_, and *d*_HH_; the intersection of these two spheres with the 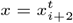 plane yields two candidates 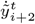 and 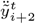 (Fig. 2c). MDCompress spends one bit to store which one is more accurate. With 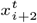 and 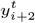 estimating 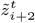 is straightforward.

If, for any reason, the estimations failed (e.g., due to errors in simulation, the spheres do not intersect), MDCompress uses the oxygen atom location as the prediction for the hydrogen atoms.

#### Predicting *MOL* positions

Predicting atom positions in the *MOL* segment is more complex. For the first three atoms, we use: at 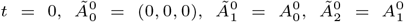 for 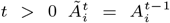. For each later atom *i*, let *p*(*i*), *q*(*i*), and *r*(*i*) denote the indices of its three reference atoms from the preprocessed model, listed in order of increasing average distance from atom *i*.

In compression levels 1, 2, and 3, we estimate 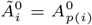,*i* ≥ 3 for the anchor frames. For the delta frames, we calculate the vector of movement of *p*(*i*)-th atom from frame *t* − 1 to frame *t*, i.e, 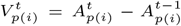 and use it for predicting the *i*-th atom coordinates i.e., 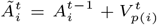 In higher compression levels, for predicting anchor atoms, we follow a method similar to that in [4]. We take 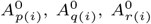 and the distances between the *i*-th atom and *p*(*i*)-th, *q*(*i*)-th, *r*(*i*)-th from the model. This 3D triangulation enables the identification of 2 compatible predicted positions 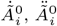 that are at the same distances from the reference points as in the model (Fig. 3a). In compression level 4, we use the better one (marking the choice with 1 bit). In compression level 5, the reference atoms define a coordinate system, and we check if the current atom is above the plane defined by the three reference atoms. We compare this information with that stored in the model to predict which of the two candidates for 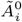 is more likely, saving 1 bit of storage.

**Fig. 3.**
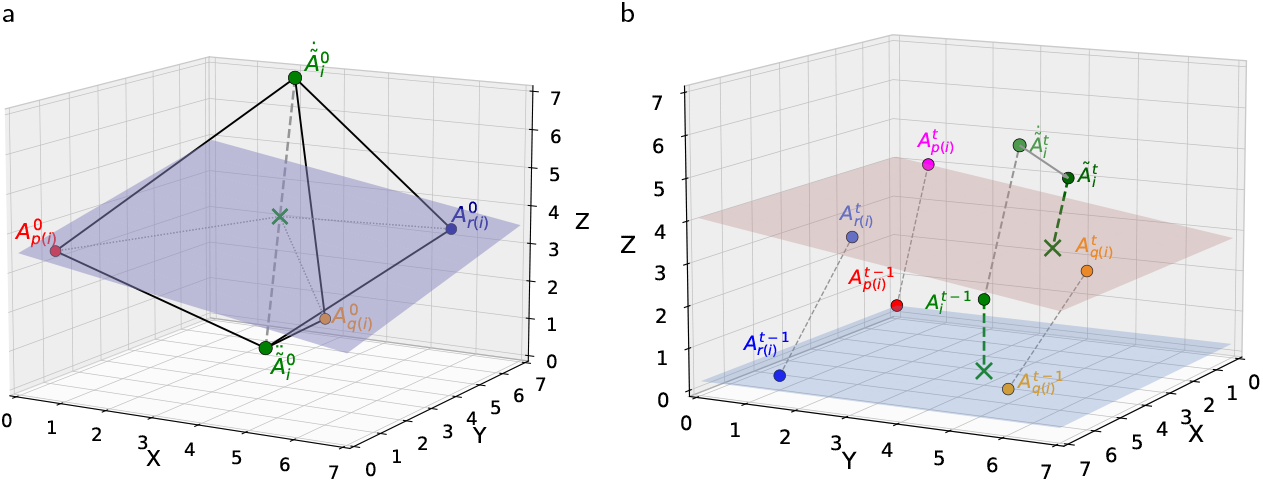
Prediction of molecule atoms for large molecule segments. (a) Prediction of coordinates for atom *A*_*i*_ in an anchor frame. Considering the positions of the three reference atoms (*A*_*p*(*i*)_,*A*_*q*(*i*)_, and *A*_*r*(*i*)_), we compute the two points 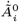 and 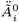 that match median distance of each reference atom to *A*_*i*_, as stored in the model. (b) Prediction of atom coordinates in delta frames, based on translation and rotation of a plane containing the reference atoms. Reference atoms from the *v*-th frame define a coordinate system *S*^*v*^, for any *v*. The point 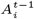 belongs to the coordinate system *S*^*t*−1^. We use translations and rotations of the coordinate system *S*^*t*−1^ (together with 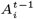) into *S*^*t*^. The position of the transformed 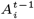 point is our prediction of 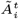 An intermediate position 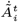 of 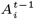 after translation is also presented.

For the delta frames, 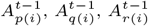 define a coordinate system *S*^*t*−1^, while 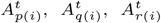,define a coordinate system *S*^*t*^. The point 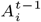 belongs to the coordinate system *S*^*t*−1^. We calculate coordinates 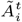 in the coordinate system *S*^*t*^. For that, we use a translation using vector 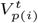, obtaining the point 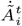 Then, we perform rotations to obtain the final prediction 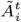 (Fig. 3b). In the case that one of the planes cannot be determined due to reference atom co-linearity, we use a vector of movement of the *p*(*i*)-th atom to estimate 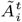.

The metadata, e.g., box of a frame, is stored in a separate container.

### Tuning compression

Various facets of the MDCompress pipeline, optionally controlled by user-selected parameters, affect compression ratio and (de)compression speed.

Considering more initial frames to determine model parameters can improve the model’s accuracy, but more than 100 frames (default) rarely helps.

The use of larger batches typically yields better compression ratios, as delta frames can be compressed more effectively than anchor frames. This has a small impact on the time required to decompress the whole trajectory. Importantly, the compression format can support efficient access to a sparse subset of trajectory frames; in case of such a request, smaller batches are better: to access each selected frame *t*, MDCompress needs to decompress just the anchor *t*^*′*^ preceding *t* and all delta frames between *t*^*′*^ and *t*.

The MDC format also enables efficient retrieval of the complete trajectories of a sparse subset of few atoms, since segments are internally split into smaller parts (sub-segments) containing 100 atoms (user-defined parameter). Thanks to this, when some atom coordinates are necessary, we only need to decompress a small sub-segment rather than a much bigger segment (cf. Fig. 1b, zoomed part). If such functionality is unnecessary, the user can turn subsegments off, gaining a bit in compression ratio.

### Implementation

The main utility, MDCompress, is provided as a command-line tool. Figure 4a shows the architecture of the compression mode. The input can be a trajectory file in XTC, TRR, TNG, NC, or GRO formats (for parsing we use chemfiles library [23]). In decompression, one can choose among XTC, TRR, NC, GRO formats. Each yellow box represents a single thread. The Reader loads the frames one by one, gathers them in batches of user-specified size, and stores them in a bounded queue. The Splitter thread splits each batch into subsegments, which are sent to a queue as input to the Worker threads that perform the core compression. The compressed blocks are passed to the final priority queue. Then, the Storer thread gathers the blocks in a deterministic order (to preserve archive binary compatibility independent on the number of threads used for compression) and stores them in a final archive file. MDCompress launches more threads than the number specified by the user. Nevertheless, the number of simultaneously working threads never breaks this limit.

**Fig. 4.**
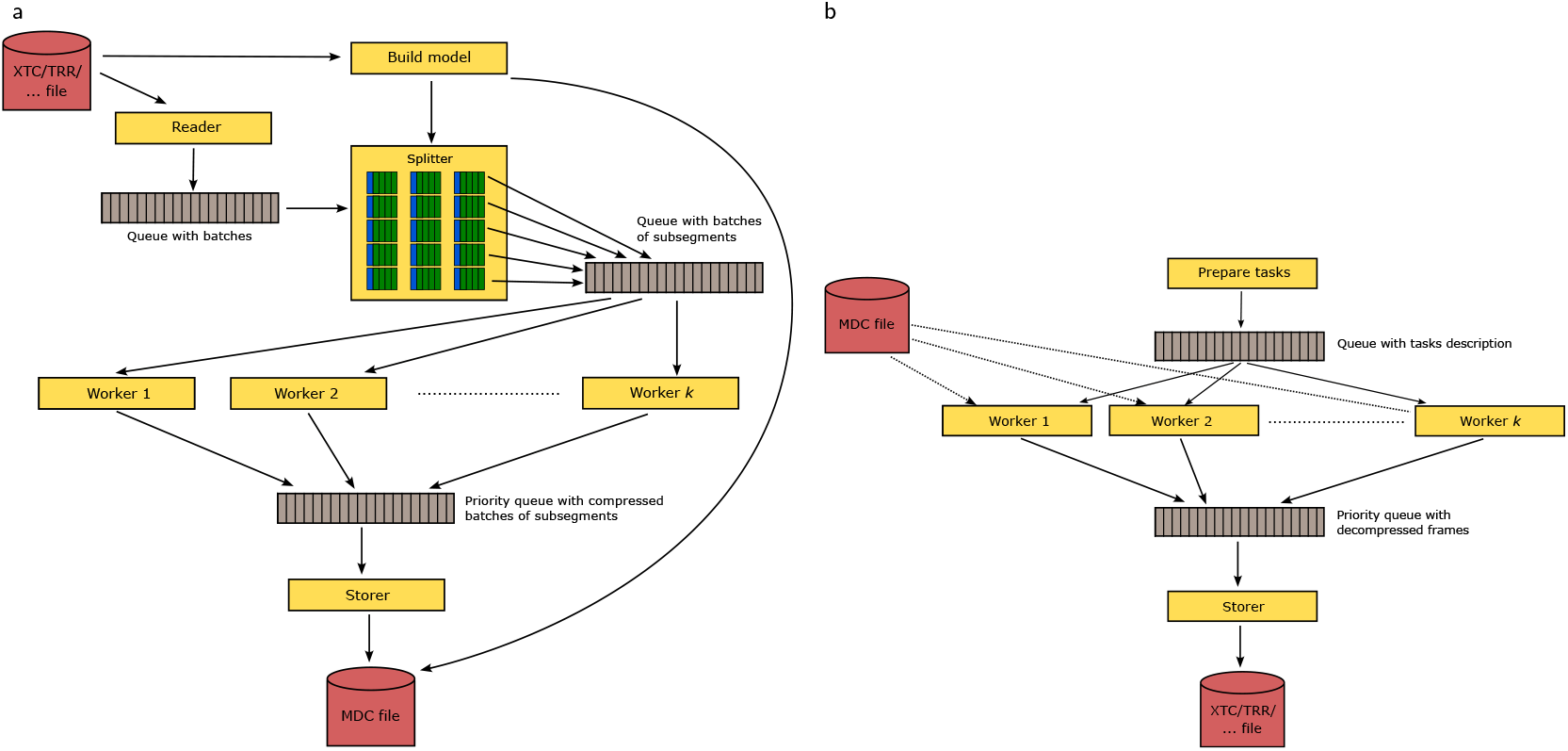
Architecture of MDCompress tool. The yellow boxes represent threads. The gray boxes represent queues (FIFO by default). The red boxes represent files. (a) Data flow in compression. (b) Data flow in decompression and access queries.

Figure 4b shows the decompression and query modes. The user’s request is initially analyzed, and a description of the necessary decompression tasks is prepared. Then, the maximum allowed number of Worker threads is launched. Each of them takes a single task and accesses the MDC file for the input data. The results are sent to a priority queue, from which the Storer thread takes them in the correct order to save in one of the allowed output formats, e.g., XTC, TRR. In this mode, we also activate only the maximum allowed number of simultaneously operating threads.

### Archive format

The MDC archive is a container database, similar to that used in [4]. Technically, the database comprises *streams*, each containing any number of *parts*. A *part* is a block of bytes with some small metadata (single 64-bit integer). Each part of each stream can be accessed in a random-access manner. The format is easily extendable, so any streams can be added to the archive at any time, and the file will still remain valid.

At the moment, we use three streams for the metadata: frame IDs, segment descriptions, and anchor IDs (the format allows for batches of variable size; however, this is not implemented yet in the compressor). We also use a family of streams to store models (such as *d*_OH_, *d*_HH_, and reference atoms) for each subsegment. Optionally, there is also a stream for storing the topology file provided by the user. Finally, there is a family of streams for atom coordinates within subsegments. In the future, we plan to add the other streams, e.g., for forces and velocities.

### Portability

MDCompress is implemented in C++20 programming language and produces a standalone executable for compressing and decompressing molecular trajectories. For easy adoption, the package also provides thread-safe decompression libraries for C, C++, Python, Rust, and WebAssembly. The libraries share a largely language-independent architecture. Implementation details for using the libraries are provided in the user guide.

For maximum portability, all coordinate calculations use our custom fixed-precision floating-point numbers.

## Experimental results

For the experiments, we primarily used 30 simulations of varying protein sequences, downloaded from MDRepo [11]; these simulations are derived from the ATLAS database [7] and the full simulations range in size from 1.9 GB to 13.8GB in uncompressed TRR format. We also evaluated compression of two larger simulations that we recently contributed to MDRepo: MDR00020616 (a protein kinase in complex with a small molecule ligand, 96.1 GB) and MDR00004434 (a Nicotinamide phosphoribosyltransferase in complex with a ligand, 60.7 GB).

The repository MDRepo makes two representations of each simulation available: (1) the *full* trajectory containing all molecules in the simulation and (2) a *minimal* trajectory from which bulk solvent (water) and ions have been removed. In most simulations, water molecules make up ~90% of the atoms, so minimal trajectory files are typically one-tenth the size of the corresponding full trajectory. Minimal trajectories are useful in cases when solvent and ion coordinates are not required, such as in training AI models for protein interaction or structure–function prediction, and are therefore expected to dominate large-scale data transfers in practice. Since MDCompress is more effective in compressing macromolecules than solvent and ion atoms, it should produce higher compression ratios for minimal trajectories. To evaluate this effect, we report performance benchmarks on both *full* and *minimal* trajectories.

We also explored the impact of varying time-steps between frames by considering a simulation (MDR00020615) sampled at an extremely dense sampling timestep of 0.1 ps, and producing downsampled versions of the simulation with timesteps of 0.3 ps, 1 ps, 3 ps, and 10 ps.The characteristics of these datasets are provided in Supplementary Worksheet.

The experiments were made on an AMD 3995WX CPU-based machine with 1 TiB RAM. To minimize the impact of variable I/O performance, the data were stored on an NVMe drive. Unless otherwise stated, MDCompress was run with 8 threads; other tools do not support multithreading, so were run with 1 thread.

In all tests, we used default values (batch size = 20, subsegment size = 100, compression level = 2, no. frames for model = 100) for all parameters except where otherwise noted.

### Examining impact of parameter values on performance

At higher compression level settings, more calculations are used to better predict the positions of molecules at higher levels. As expected, running MDCompress with a higher compression level leads to increased compression ratio (defined as the size of the TRR file divided by the size of the compressed file) (Fig. 5a), while the speed of both compression and decompression of the whole dataset decreases (Fig. 5b). Levels 2 or 4 seem the most practical, depending on the users’ needs.

**Fig. 5.**
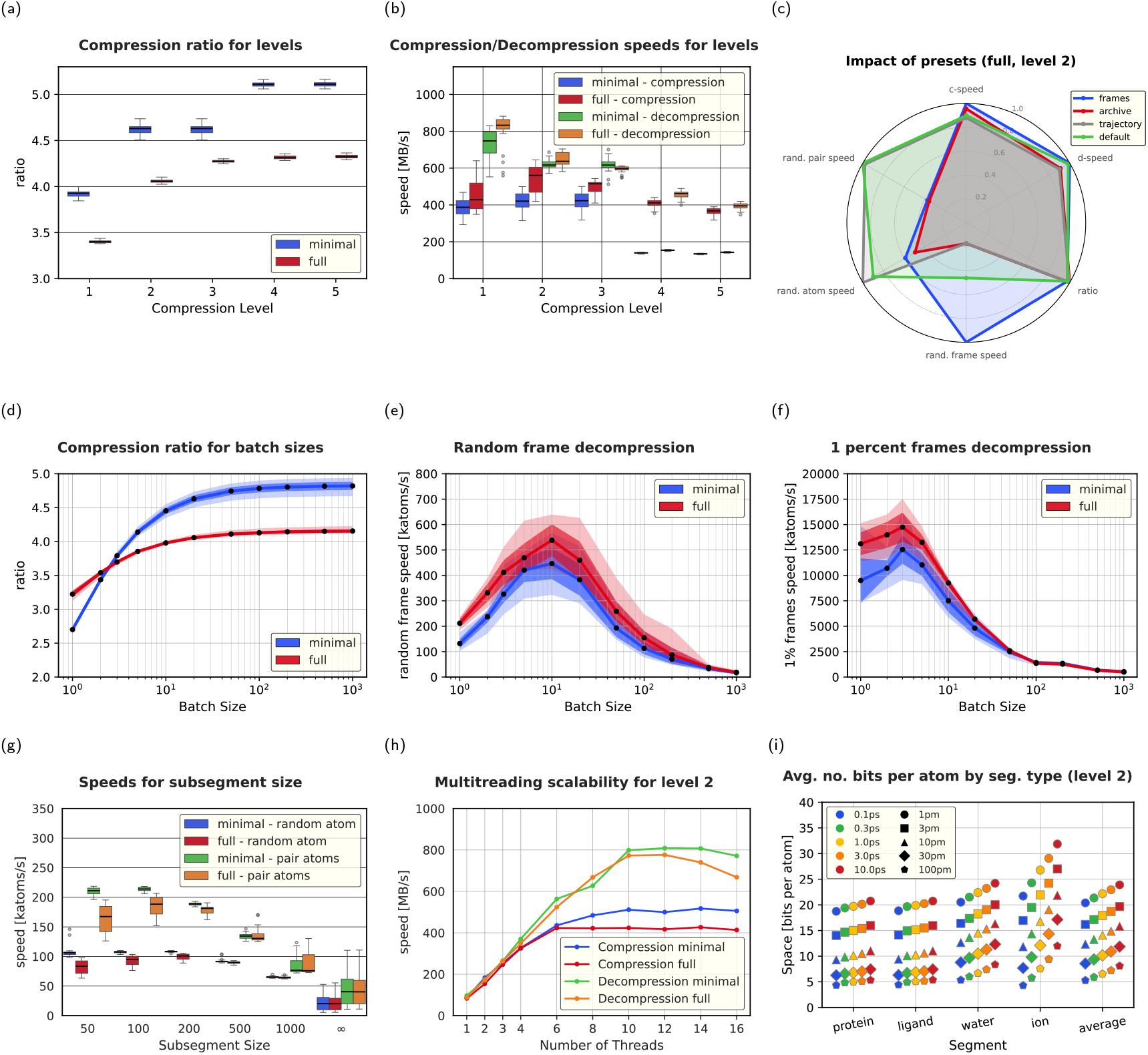
Impact of parameters on compression speed and ratio, various types of queries. Some charts show two groups: *full* (all atoms, including water) and *minimal* (water and ion atoms removed). All charts (except for **(h)** and **(i)**) show the results for a dataset of 30 simulations taken from MDRepo (see Supplementary Worksheet for their characteristics). Median, std. dev, and min/max values are shown on box plots and range plots. **(a)** Compression ratio (uncompressed size / compressed size) for various compression levels. **(b)** Compression and decompression speed for various compression levels. Impact of presets on compression ratio, (de)compression speeds, and three types of queries. Compression level 2 and a trajectory dataset are used. Compression ratio for various batch sizes. **(e)** Decompression rate when extracting a random frame, for various batch sizes. **(f)** Decompression rate for extracting 1 percent frames evenly spread across the trajectory, for various batch sizes. **(g)** Decompression speed for the whole trajectory of one or two atoms. **(h)** Compression speed in terms of number of threads, using the dataset MDR00004144. **(i)** No. of bits necessary to store atoms from: the protein, the small molecule ligand, all water molecules, all ions in the MDR00020615 dataset. Various frame sampling steps and resolutions are presented.

**Fig. 6.**
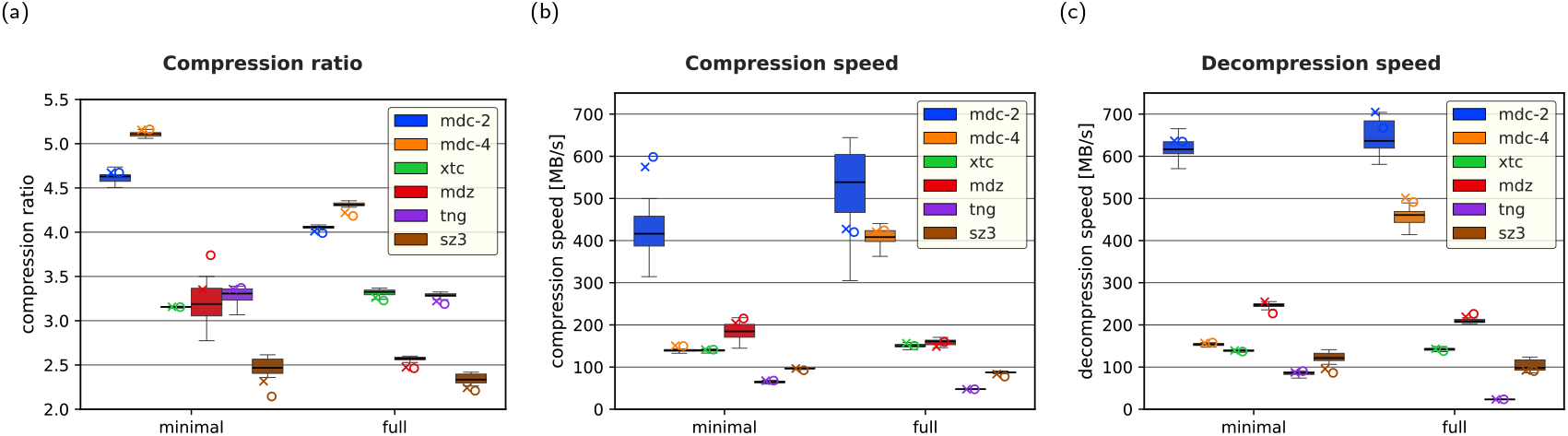
Comparison of the examined tools. Each chart shows two groups: minimal (only molecule atoms) and trajectory (all atoms, including water). MDCompress was examined for compression levels 2 and 4 (in both cases, the default preset). Three series are presented: box plots for a 30-file ATLAS dataset. Median, std. dev, and min/max values are shown. The ‘X’ marker is for simulation MDR00004434, and the ‘O’ marker is for simulation MDR00020616. **(a)** Compression ratios. **(b)** Compression speed. **(c)** Decompression speed.

Using large batch sizes means that very few frames are anchors and many are delta-encoded, which improves compression. This is reflected in Fig. 5d, which shows that compression ratios grow with increasing batch size, approaching a maximum compression ratio that matches the ratio observed in delta frames. On the other hand, when requesting a subset of frames from a file, it is preferable to have smaller batches, because decompressing each target frame *t* requires analyzing the preceding anchor frame *a* and all frames between *a* and *t*. We evaluated decompression speed (measured as the number of decompressed atoms per second) when selecting a single random frame (Fig. 5e) and when sampling 1% of frames with a constant stride (Fig. 5f). Batch sizes of around 5–10 frames yield maximum decompression speed – larger batches require that an increasingly large number of frames must be wastefully decompressed, while very small batches require substantial overhead just to read and deserialize the metadata containing the starting positions of the very large number of batches.

To enable the user to adapt to special needs easily, we provide four presets: **default**—suggested for most scenarios (batch size = 20, subsegment size = 100), **archive**—for streaming access to whole frames or their segments (batch size = 100, subsegment size = 1000), **trajectory**—for quick access to selected atom trajectories (batch size = 100, subsegment size = 100), **frames**—for quick access to selected frames (batch size = 10, subsegment size = 1000). The comparison of the performance of these presets is presented (Fig. 5c).

By breaking segments into subsegments, MDCompress supports faster decompression for a small sample of all atoms in the system: because decompression must scan linearly through all atoms in the (sub)segment, a request to track a single atom through the simulation will benefit from decompressing only a few nearby atoms. The gain in compression ratio for larger subsegments is small (data not shown), but the speed of decompression of trajectories of single atoms or pairs of atoms differs significantly (Fig. 5g). This motivates the decision to use a small subsegment size (100) as the default.

### Multithreading performance

MDCompress is a multithreaded application, so we also evaluated its scalability. Fig. 5h shows strong scaling performance with compression level 2: MDCompress obtains compression speeds more than 400 MB/s and decompression speeds over 700 MB/s.

### Impact of the time step between sampled frames

In MD simulations, atomic positions are typically updated every 2–4 femtoseconds (the integration timestep), and a trajectory file is created by periodically saving snapshots; snapshot intervals usually range from 1ps to 1ns, depending on simulation length and whether analyses focus on fast-moving (loop or side-chain) dynamics or slow conformational changes.

In MDCompress, atom position predictions in each delta frame *t* depend in part on the position of atoms in frame *t* − 1. The accuracy of these predictions, and thus, the compressibility of the trajectory, is expected to be higher when the displacement of atom *A*_*i*_ between frames *t* − 1 and *t* is small. Thus, smaller sampling timesteps should allow for better compression. We explored this expectation by considering a simulation of a kinase protein in complex with a small molecule drug (MDRrepo entry MDR00020615) that was sampled at a 0.1 ps time step, and producing a downsampled version of the simulation by taking every 3rd, 10th, 30th, and 100th frame (equivalent to 0.3 ps, 1 ps, 3 ps, and 10 ps sampling frequencies, respectively). The results seen in Fig. 5i confirm the expectations that prediction performs better for 0.1 ps than for larger sampling steps. Moreover, the prediction for large or medium-sized molecules (protein and small molecule) is better than for water molecules; this is expected, since atom positions in large molecules can be more tightly constrained based on information from nearby reference atoms. Unsurprisingly, the hardest to compress are ion segments, which receive no positional information from neighboring atoms. The average space necessary per single atom is between 20 and 24 bits, a roughly 4.0–4.8 compression ratio relative to 96 bits in TRR format (without compression). MDCompress allows various precisions of storage of coordinates. We used several values: 1 pm (default), 3 pm, 10 pm, 30 pm, and 100 pm. The number of bits necessary to store each atom’s coordinates are also presented in the same figure. As can be observed, a remarkable reduction of space is possible if the user can sacrifice the precision.

### Comparison to other compression tools

Finally, we evaluated MDCompress against existing tools. We used XTC and TNG file formats as they are state-of-the-art compressed formats used in practice. Moreover, we included SZ3 [20] and MDZ [21] as the leading research methods proposed recently. Both SZ3 and MDZ are provided only as experimental codes, so they are not ready to be used in practice.

Nevertheless, it is interesting to see how well they perform. MDCompress was run in two modes: default (mdc-4 label) and with compression level 2 (mdc-2 label). In terms of compression ratio, MDCompress is a clear winner (Fig. 1a). It offers 15– 37% smaller archives than the competing formats XTC and TNG. The advantage is more significant for a dataset without water molecules. This is expected, as predicting the coordinates of water molecules is less successful than predicting those of atoms in longer molecules. Compression and decompression speeds (Fig. 1b–c) are comparable or better than those of the competing methods. Nevertheless, it should be emphasized that only MDCompress implements parallel processing due to the MDC format design.

## Conclusion

We have introduced MDCompress, a new software tool for compressing molecular dynamics trajectories that achieves 15–37% smaller file sizes than the widely-used XTC format while maintaining fast, multithreaded compression and decompression. The MDC format’s container-based architecture enables efficient random-access queries, useful for extracting sparse frame samplings or tracking individual atom trajectories. These capabilities are designed to meet challenges that arise as molecular dynamics datasets grow in scale and are increasingly reused for downstream analysis and AI training. At that scale, raw data volumes quickly become unwieldy, and the ability to store trajectories compactly, move them efficiently between sites, and extract only the subsets needed for a specific analysis becomes not just a convenience but a necessity for enabling real workflows. The MDCompress software is freely available under a BSD 3-clause license, with thread-safe decompression libraries provided for C, C++, Python, Rust, and WebAssembly.

Several avenues for future development remain. While our benchmarks focused on protein simulations, MDCompress handles arbitrary molecules with six or more atoms, including nucleic acids and lipids. However, solvent-specific optimizations are currently limited to water. Many other common solvents (such as methanol, glycerol, and DMSO) contain six or more atoms and can therefore be processed using the MOL segment approach, though this requires explicit specification for each molecule, and optimizes only a subset of atomic positions; meanwhile smaller solvent molecules are handled as OTH segments with baseline compression comparable to XTC. We aim to expand optimized solvent handling in future releases.

## Supporting information

Supplementary Worksheet

## Author contributions statement

S.D. and T.J.W. designated functional requirements and specifications. S.D. and M.K. designed data structures and algorithms. S.D. and M.K. implemented the software. A.R. and T.J.W. prepared data samples (gathering existing simulations, preparing new simulations, creating data handling scripts). All authors conceived the experiments. M.K. conducted the experiments and analysed results. S.D. and M.K. prepared figures. T.J.W. and S.D. wrote the first draft of the manuscript and all authors revised aspects of the manuscript.

## Data availability

All data used in validation is available from https://mdrepo.org, using the unique identifiers listed in Supplementary Worksheet.

