## Supplementary Material for "MDCompress: better, faster compression of molecular dynamics simulation trajectories"

### Supplementary data

#### Command lines used in the experiments

Times and memory use were measured by `\usr\bin\time -v` command.

All experiments used TRR files as input. Because MDRepo stores data in XTC format, the data was compressed to MDC using mdcompress and then decompressed to TRR.

##### XTC

GROMACS 2025.1 package was used to evaluate XTC compression.

- Compression:
  ```
  /usr/bin/time -v gmx trjconv -f <input.trr> -o <output.xtc>
  ```
- Decompression:
  ```
  gmx trjconv -f <input.xtc> -o <output.trr>
  ```

##### TNG

GROMACS 2025.1 package was used to evaluate TNG compression.

- Compression:
  ```
  /usr/bin/time -v bash -c 'echo 0 | gmx trjconv -f "$1" -o "$1.xtc" -s "$2"' _ "<input.trr>" "<topology.pdb>"
  /usr/bin/time -v bash -c 'echo 0 | gmx trjconv -f "$1.xtc" -o "$1.tng" -s "$2" -round' _ "<input.trr>" "<topology.pdb>"
  ```
  Unfortunately, we were not able to compress directly from TRR to TNG with rounding to 1 pm. That is why we went through XTC temporary file.
- Decompression:
  ```
  /usr/bin/time -v gmx trjconv -f "<input.tng>" -o "<outut.trr>"
  ```

### SZ3

SZ3 downloaded from `https://github.com/szcompressor/SZ3` (commit: `e8a6b1569067abdd6b7d4276e91eced115be4f14`) was used for evaluation. This is an experimental code. Thus, we needed to convert TRR input file into a plan file with float values (32-bit).

- Compression:
  ```
  /usr/bin/time -v ./sz3 -i <input.fp32> -z <output.sz3> -f -M ABS 0.005 -3 <no_frames> <no_atoms> 3
  ```
  The absolute max. error value used was to have the same precision as for XTC, TNG, MDCompress.
- Decompression:
  ```
  /usr/bin/time -v ./sz3 -z <input.sz3> -o <output.fp32> -f -M ABS 0.005 -3 <no_frames> <no_atoms> 3
  ```

##### MDZ

SZ3 downloaded from `https://github.com/szcompressor/SZ3` (commit: `e8a6b1569067abdd6b7d4276e91eced115be4f14`) was used for evaluation. One of its subtools implements MDZ compression. This is an experimental code. Thus, we needed to convert TRR input file into a plan file with float values (32-bit). Moreover, the tool only does compression and decompression in memory, so no file is produced. The results are taken from the output as MDZ reports the compression ratio. The tool often crashes and reports a compression ratio 1.0. To make it work, we fixed two bugs. First, we read the compressed file size from the function doing the compression (it was ignored). Second, we changed the allocated space for internal ZSTD compression (the originally used value was too small).

- Compression and decompression:
  ```
  /usr/bin/time -v ./mdz <input.fp32> -3 <no_frames> <no_atoms> 3 -a 0.005 20 2
  ```
  The absolute max. error value used was to have the same precision as for XTC, TNG, MDCompress. The batch size used (20) was selected to meet the default batch size of MDCompress.

##### MDCompress

To test various batch sizes:

```
/usr/bin/time -v ./mdcompress compress -b <batch_size> -i <input.trr> --topology <topology_path> -o <output.mdc>
```

To test various compression levels:

```
/usr/bin/time -v ./mdcompress compress -l <level> -i <input.trr> --topology <topology_path> -o <output.mdc>
```

To test various subsegment sizes:

```
/usr/bin/time -v ./mdcompress compress --subsegment-size <subsegment_size> -i <input.trr> --topology <topology_path>
-o <output.mdc>
```

To test presets:

```
/usr/bin/time -v ./mdcompress compress -l <level> --preset <preset> -i <input.trr> --topology <topology_path> -o <output.mdc>
```

All the above command lines were extended with `--only-mol` flag for the minimal files.

To test full decompression:

```
/usr/bin/time -v ./mdcompress select -i <input.mdc> -o <output.trr>
```

To test 1% of frames decompression:

```
/usr/bin/time -v ./mdcompress select --fr <start_frame_id> <no_frames - 1> --stride 100 -i <input.mdc> -o <output.trr>
```

In this case, start_frame_id was set to $\lfloor \texttt{batch\_size}/2 \rfloor$.

To test decompression of a single random frame:

```
/usr/bin/time -v ./mdcompress select --fid <random_frame_id> -i <input.mdc> -o <output.trr>
```

To test decompression of a single random atom:

```
/usr/bin/time -v ./mdcompress select --atoms <random_atom_id> -i <input.mdc> -o <output.trr>
```

To test decompression of a pair of random atoms:

```
/usr/bin/time -v ./mdcompress select --atoms <first_random_atom_id>,<second_random_atom_id> -i <input.mdc> -o <output.trr>
```

To test decompression of a pair of close random atoms:

```
/usr/bin/time -v ./mdcompress select --atoms <first_random_atom_id>,<second_random_atom_id> -i <input.mdc> -o <output.trr>
```

With the additional condition that $\texttt{second\_random\_atom\_id} - \texttt{first\_random\_atom\_id} > 50$.

In the case of random frame, random atom, pair of random atoms and pair of random atoms at maximum distance 50, tests were run 21 times and the median value was reported. In the case of random atoms, only non-water atoms were considered.

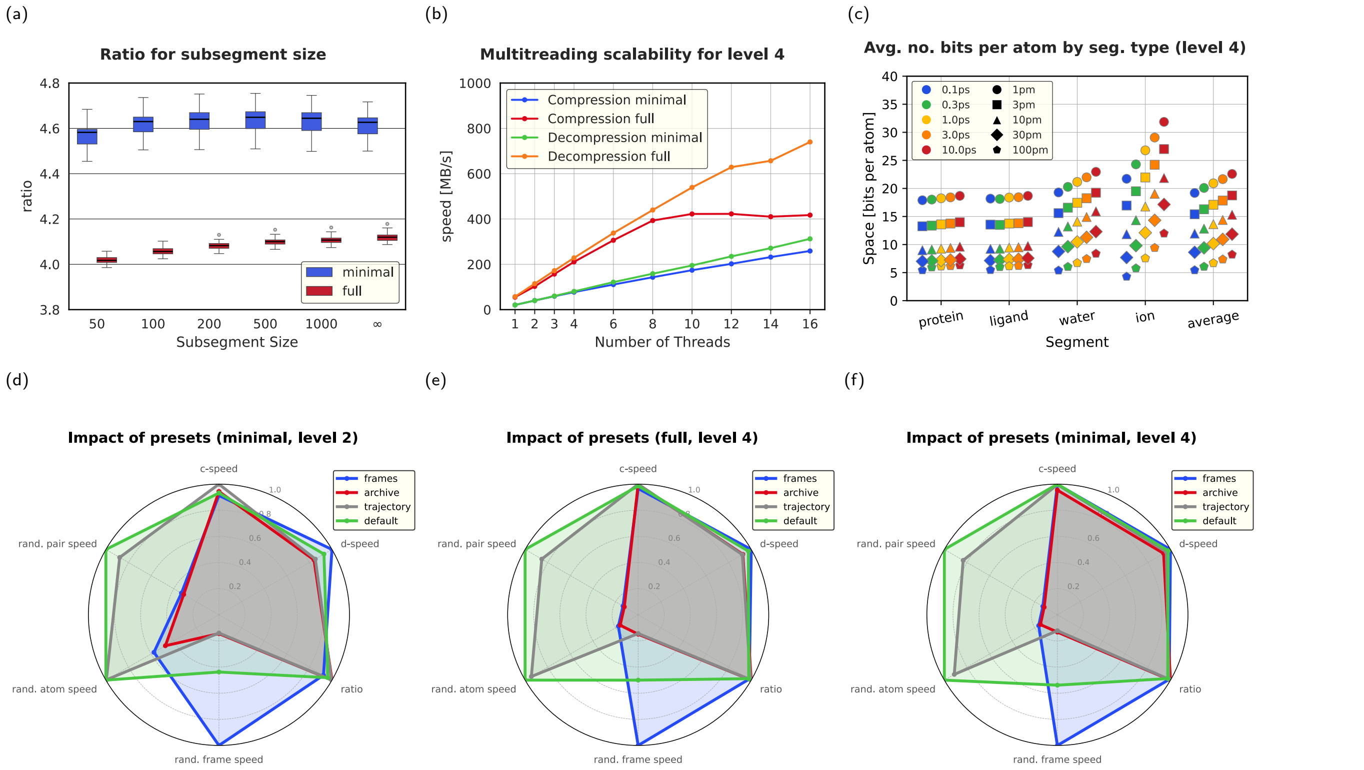


**Fig. 1.** **(a)** Compression ratio for various subsegment sizes. **(b)** Decompression speed in terms of number of threads for compression level 4. **(c)** No. of bits necessary to store atoms from: the protein, the small molecule ligand, all water molecules, all ions in the MDR00020615 dataset for compression level 4. **(d)** Impact of presets on compression ratio, (de)compression speeds, and three types of queries. Compression level 2 and a minimal dataset are used. **(e)** Impact of presets on compression ratio, (de)compression speeds, and three types of queries. Compression level 4 and a full dataset are used. **(f)** Impact of presets on compression ratio, (de)compression speeds, and three types of queries. Compression level 4 and a minimal dataset are used.
